# Transcriptomic Profiling of Thyroid Eye Disease Orbital Fat Demonstrates Differences in Adipogenicity and IGF-1R Pathway

**DOI:** 10.1101/2024.04.19.590238

**Authors:** Dong Won Kim, Soohyun Kim, Jeong Han, Karan Belday, Emily Li, Nicholas Mahoney, Seth Blackshaw, Fatemeh Rajaii

## Abstract

Despite recent advances in the treatment of thyroid eye disease (TED), significant gaps remain in our understanding of the underlying molecular mechanisms, particularly concerning the insulin-like growth factor-1 receptor (IGF-1R) pathway. To dissect the pathophysiology of TED, we utilized single nucleus RNA-Seq to analyze orbital fat specimens from both TED patients and matched controls. The analysis demonstrated a marked increase in the proportion of fibroblasts transitioning to adipogenesis in the orbital fat of TED patients compared to controls. This was associated with diverse alterations in immune cell composition. Significant alterations in the IGF-1R signaling pathway were noted between TED specimens and controls, indicating a potential pathological mechanism driven by IGF-1R signaling abnormalities. Additionally, our data show that linsitinib, a small molecule inhibitor of IGF-1R, effectively reduces adipogenesis in TED orbital fibroblasts *in vitro*, suggesting its potential utility as a therapeutic agent. Our findings reveal that beyond immune dysfunction, abnormal IGF-1R signaling leading to enhanced adipogenesis is a crucial pathogenic mechanism in TED.

## Introduction

Autoimmune thyroid disease is the most prevalent organ-specific autoimmune disease, affecting 2-5% of the global population. Approximately 25-40% of patients with autoimmune thyroid disease eventually develop thyroid-related eye disease (TED), a condition that can lead to disfigurement and vision loss (1–4). Although the pathophysiology of TED is not entirely understood, it is thought that an immune response targeting autoantigens present on thyrocytes and orbital fibroblasts, the thyroid stimulating hormone receptor (TSHR) and/or the insulin-like growth factor-1 receptor (IGF-1R), induces a cascade of inflammatory cytokines (5–15). This process increases orbital soft tissue volume through mechanisms such as glycosaminoglycan production, fibrosis, and adipogenesis, all of which have been demonstrated *in vitro* (12, 13, 15). The expansion of orbital soft tissues, responsible for proptosis, diplopia, pain, and vision loss in TED, is thought to be the final step of inflammatory cascades that result in fibrosis and adipogenesis.

IGF-1 enhances the expression of TSHR in orbital fibroblasts, and treatment with the monoclonal anti-IGF-1R antibody teprotumumab reduces TSHR and IGF-1R expression (16). Moreover, teprotumumab blocks the induction of inflammatory cytokines interleukin 6 (IL-6), and IL-8 in orbital fibroblasts (17). Single nucleus RNA sequencing (snRNA-Seq) has identified that during the early stages of adipogenesis, orbital fibroblasts upregulate the expression of both IGF-1 and IGF-1R *in vitro* (18).

In this study, we aimed to demonstrate the role of adipogenesis in TED *in vivo*. We utilized snRNA-Seq to compare gene expression profiles in retrobulbar fat from control patients undergoing orbital surgery and compared it to that of TED patients undergoing orbital decompression. Our findings not only confirmed an increase in adipocytes but also highlighted an increased presence of fibroblasts undergoing adipogenesis in the orbital fat of TED patients. Furthermore, we observed disruptions in the IGF pathway, including IGF-1R signaling in both the orbital fibroblasts and fat of TED patients. To investigate the function of the IGF-1R signaling pathway in orbital adipogenesis, we employed linsitinib, a small molecule antagonist of IGF-1R. Our *in vitro* studies demonstrated that blocking IGF1-R signaling reduces adipogenesis in TED orbital fibroblasts, providing new insights into the pathophysiological mechanisms of TED and identifying potential therapeutic targets.

## Methods

### Cell culture

Orbital fibroblast cell lines were derived from the retrobulbar fat of TED and control patients, following a protocol approved by the Johns Hopkins University Institutional Review Board and following the tenets of the Declaration of Helsinki as previously described (19, 20). Briefly, tissue explants were obtained from patients undergoing surgical decompression for TED or excision of prolapsed orbital fat, which has been shown to be retrobulbar fat, in control patients (19). TED patients were inactive, with a clinical activity score (CAS) less than 4, at the time of surgery (21). The explants were placed on the bottom of culture plates and covered with Eagle’s medium containing 10% fetal bovine serum (FBS), antibiotics, and glutamine, and then incubated at 37^○^C, 5% CO_2_, in a humidified environment. The culture media was refreshed weekly until cell confluence was achieved. The resulting fibroblast monolayers were passaged serially by treatment with TrypLE (#12604039, ThermoFisher Scientific, Waltham, MA), and maintained in liquid nitrogen storage for future use.

For adipogenesis induction, orbital fibroblasts were induced to undergo adipogenesis as previously described (20). Briefly, fibroblasts between passages 4 and 8 were seeded on plastic tissue culture plates until near-confluence in DMEM containing 10% FBS and antibiotics. Adipogenic differentiation was initiated by treating the cells with adipogenic medium composed of DMEM:F-10 (1:1, #11320033, ThermoFisher Scientific, Waltham, MA) supplemented with 3% FBS, 100 nmol/liter insulin (#12585014, Thermo Fisher Scientific, Waltham, MA), 1 μmol/liter dexamethasone (#D4902, Sigma, St. Louis, MO), and for the first week only, 1 μmol/liter rosiglitazone (#R2408, Sigma, St. Louis, MO) and 0.2 mmol/liter IBMX (#13630S, Cell Signaling Technology, Danvers, MA). Linsitinib (#T69071, TargetMol, Wellesley Hills, MA) was added to the adipogenic media at the indicated concentration during the first week only. The medium was replaced every other day for the first week and the culture was continued in this medium for a total of 9 days. Control cultures were treated with DMEM:F-10 (1:1), supplemented with 3% FBS. Differentiation was observed using a Nikon microscope (Japan), with each time point replicated in triplicate.

### Quantification of adipocytes

On days 0, 5, and 9 of the experiment, cells were washed with PBS followed by fixation with 4% paraformaldehyde in PBS for 10 minutes at room temperature. Cells were then washed with PBS or stored in PBS at 4^○^C. For adipocyte staining, cells were first washed with 60% isopropanol. Staining was then performed with freshly prepared 0.3% (w/v) Oil red O (Sigma, St. Louis, MO) at room temperature for 15 minutes. After the staining process, cells were rinsed with 60% isopropanol and counterstained with hematoxylin (Sigma, St. Louis, MO) for 5 minutes, followed by a wash with PBS. Cells were imaged by a masked study team member using a Keyence BZ-X710 microscope (Keyence, Japan). For each treatment replicate, five high-power images were captured. Cell counts and lipid vacuole measurements were performed by a masked study team member using ImageJ (22). Adipocytes were identified as Oil Red O-positive cells, and nuclei in cells that did not stain with Oil Red O, indicative of fibroblasts or preadipocytes, were excluded from the adipocyte count. Data is presented as mean with standard deviation (SD). Two-way ANOVA with Tukey’s multiple comparison analysis was performed using Prism (GraphPad, Boston, MA).

### Western Blot

Cellular protein was collected in RIPA buffer (R0278, Millipore Sigma, Burlington, MA) containing ReadyShield^®^ Protease and Phosphatase Inhibitor Cocktail (PPC- 2020, Millipore Sigma, Burlington, MA) on days 0, 5 and 9 of the experiment. The protein extracts were subjected to quantification using the DC Protein Assay Reagents (5000116, Bio-Rad, Hercules, CA). The RIPA protein samples were mixed with 4X protein sample buffer (1610747, Bio-Rad, Hercules, CA) containing 5% 2- mercaptoethanol (1610710, Bio-Rad, Hercules, CA) and heated at 95°C for 5 min to denature. Proteins were loaded onto 10% bis-Tris gels and transferred to Immun-Blot PVDF membranes (1620177, Bio-Rad, Hercules, CA), which were then blocked in 5% blocking grade skim milk (170–6404, Bio-Rad, Hercules, CA). Primary antibody was used at a dilution of 1:1000. Antibodies specific for phospho IGF-1R (#3918), IGF-1R (#3027), and actin (#5125) were obtained from Cell Signal Technology (Danvers, MA). The membranes were incubated with the appropriate primary antibody overnight followed by horseradish peroxidase-conjugated anti-rabbit (#7074, Cell Signaling Technology, Danvers, MA) secondary antibody at a dilution of 1:2500 for an hour at room temperature. The signal was detected with a chemiluminescence detection system (Thermo Scientific, Waltham, MA). Blots were imaged with an iBright CL1500 Imaging System (Thermo Fisher Scientific Inc, USA), and band intensity was reported as arbitrary densitometric units using ImageJ software (National Institutes of Health, USA). Two-way ANOVA with Tukey’s multiple comparison analysis was performed using Prism (GraphPad, Boston, MA).

### snRNA-Seq

Human retrobulbar orbital fat was obtained from TED patients undergoing orbital decompression or control patients undergoing routine resection of prolapsed orbital fat (Table S1), which is intraconal fat (19), using a protocol approved by the Johns Hopkins University Institutional Review Board and following the tenets of the Declaration of Helsinki. TED patients were inactive at the time of the decision to proceed with surgery, based on a CAS of less than 4 (Table S1). Nuclei from retrobulbar orbital fat were isolated using the methods described in the 10x Genomics Sample Preparation Demonstrated Protocol (10x Genomics, Pleasanton, CA). Briefly, cells were washed with chilled PBS and lysed in chilled lysis buffer consisting of 10 mM Tris-HCl, 10 mM NaCl, 3 mM MgCl2, and 0.1% Nonidet^TM^ P40 Substitute (MP Biomedicals, Solon, OH) in nuclease-free water at 4^○^C. Cells were scraped from the plate bottom and centrifuged at 500 RCF for 5 min at 4^○^C. Cells were washed twice in nuclei wash and resuspension buffer consisting of PBS with 0.1% BSA and 0.2 U/μl RNase inhibitor (#N2615, Promega, Madison, WI). Cells were passed through a 50 μm CellTrics^TM^ filter (#NC9491906, Sysmex, Germany) and centrifuged at 500 RCF for 5 min at 4^○^C before resuspension in wash and resuspension buffer. Isolated nuclei were counted manually via hemocytometer with Trypan Blue staining, and nucleus concentration was adjusted following the 10x Genomics guideline. 10x Genomics Chromium Single Cell system (10x Genomics, CA, United States) using V3.1 chemistry per manufacturer’s instructions, generating a total of 10 libraries. A couple of biological groups were run with technical replicates (Table S1). Libraries were sequenced on Illumina NovaSeq 6000 with 500 million reads per sample. Sequencing data were pre-processed through the Cell Ranger pipeline (10x Genomics, Cellranger v5.0.0) with default parameters, using GRCh38-2020-A genome with *include-introns*.

Matrix files generated from the Cellranger run were used for subsequent analysis as described previously (18, 23). Briefly, Seurat V3 (24) was used to perform downstream analysis, only including cells with more than 500 genes, and 1000 UMI, to process control and treated samples separately. Seurat SCTransform function was used to normalize the dataset (25) and UMAP was used to reduce dimensions derived from the Harmony (26) output. Individual cell types were first identified by cross-referencing to the ASCOT database (27).

Differential gene tests on the scRNA-Seq datasets were initially performed using Seurat v3.15’*FindAllMarkers’* function (*test.use* = *“wilcox”, logfc.threshold* = *0.5, min.pct* = *0.2)*. Top differential genes (adjusted *p*-value < 0.05, fold change > 0.5) in each microglial cluster were used as input for GO pathway analysis using ClusterProfiler (v3.12.0) (28). Pseudotime analysis was performed using Monocle v3 (29) and significant genes (q value < 0.001) along pseudotime trajectory were used to run KEGG analysis as described above. Genes that were involved across KEGG signaling pathways were then used to identify signaling pathways that are specific to adipocyte differentiation.

### Sex as a biological variable

Our study examined male and female human samples (Table S1), and we did not observe any sex-dimorphic effects in this study. Details of demographics are shown in Table S1.

### Data availability

All other data reported in this paper will be shared by the authors upon request due to IRB policy against open-access sharing of human subjects data. No custom code was used in this study.

## Results

### TED human orbital tissue shows increased adipocytes

To delineate changes in the cellular composition within orbital tissue associated with TED, we utilized snRNA-Seq to analyze samples from patients diagnosed with TED in comparison to controls without TED pathology (Figure 1A, Table S1). Our analysis identified four primary cell populations present in both groups: fibroblasts, adipocytes, endothelial cells, and a diverse array of immune cells (Figure 1B). Fibroblasts were characterized by the expression of genes such as DCN, CNTN4, and ADAMTSL1 (Table S2). Adipocytes were identified through the expression of *PDE3B, LPL*, and *PLIN1* (Table S2). Endothelial and smooth cells exhibited *VWF* expression, and the immune cell population comprised multiple subsets including neutrophils, mast cells, and cells indicative of either innate (Immune cell Cluster 2, *F13A1,* and *MRC1*) or adaptive (Immune cell Cluster 1, *PTPRC* and *RIPOR2*) responses (Table S2).

**Figure 1.**
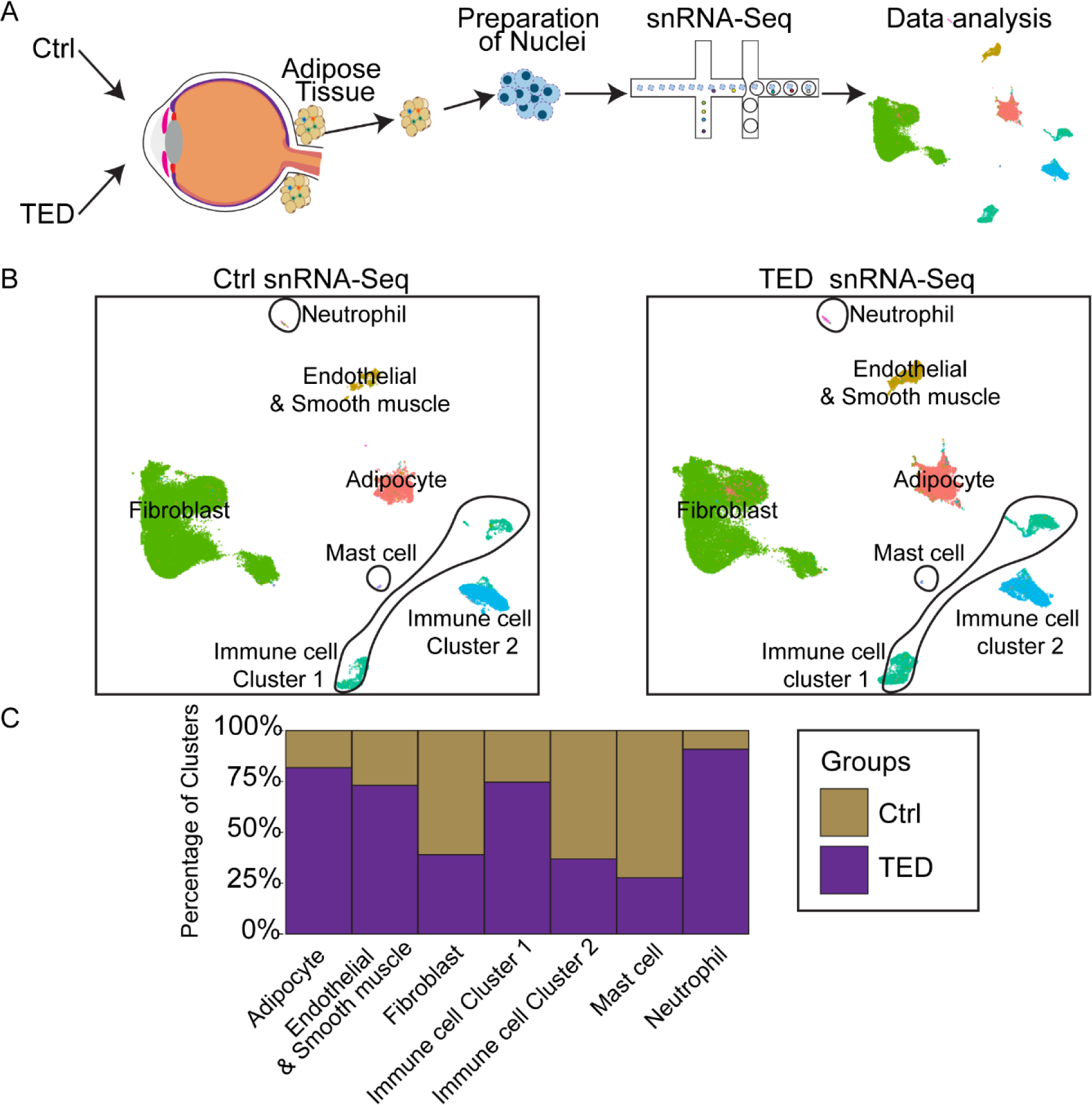
**A.** Schematic of the experimental approach. **B.** UMAP plots illustrating the distribution of cell types in control (left) and TED (right). **C.** Bar graphs comparing the distribution of control and TED groups across identified clusters in snRNA-Seq.

Notably, the TED patient group exhibited a significant increase in adipocyte numbers compared to the control group (Figure 1C, S1), consistent with the adipogenic shift previously documented in TED pathology (18). Increases in *IGF1*, *IGF1R*, and *TSHR* in TED fibroblasts compared to the controls were observed (Figure S2), aligning with previous studies (30). While increases in *IGF1* and *TSHR* were observed in TED adipocytes compared to controls, *IGF1R* levels did not differ between the groups (Figure S2). *IGF2* expression was absent in both groups (Figure S2).

Furthermore, adipogenic markers such as *PDE3B* and *PLIN1* were detected in TED fibroblasts but not in controls (Figure S3), suggesting an ongoing adipogenic transition within TED fibroblasts. Additionally, our snRNA-Seq data highlighted an increased presence of immune cells facilitating adaptive responses in TED samples, contrasting with a diminished representation of cells associated with innate immunity (Figure 1C). These observations not only validate known histopathological features in TED but also reveal new insights into the disease’s immune landscape.

### Exploring Variations in Immune Cell Distribution Between Control and TED Groups

To further explore the variations in immune cell distribution between the control and TED groups, we focused our analysis on ‘Immune Cell Clusters 1 and 2’. These clusters were characterized by gene expressions indicative of adaptive (Cluster 1) and innate (Cluster 2) immune responses (Table S2). A Gene Ontology (GO) pathway analysis of TED immune cells (Figure 2A, B, Table S3), revealed significant pathways such as ‘T cell differentiation in thymus’, and ‘positive T cell selection’. This analysis also uncovered heightened expressions of *CD74, LYZ, HLA-DBA, CD163,* and *IRAK3* in TED immune cells (Figure 2C, Table S3) - genes generally enriched in adaptive immune cells. Such findings suggest a pronounced shift toward an adaptive immune response in TED, consistent with autoimmune etiology (31, 32).

**Figure 2.**
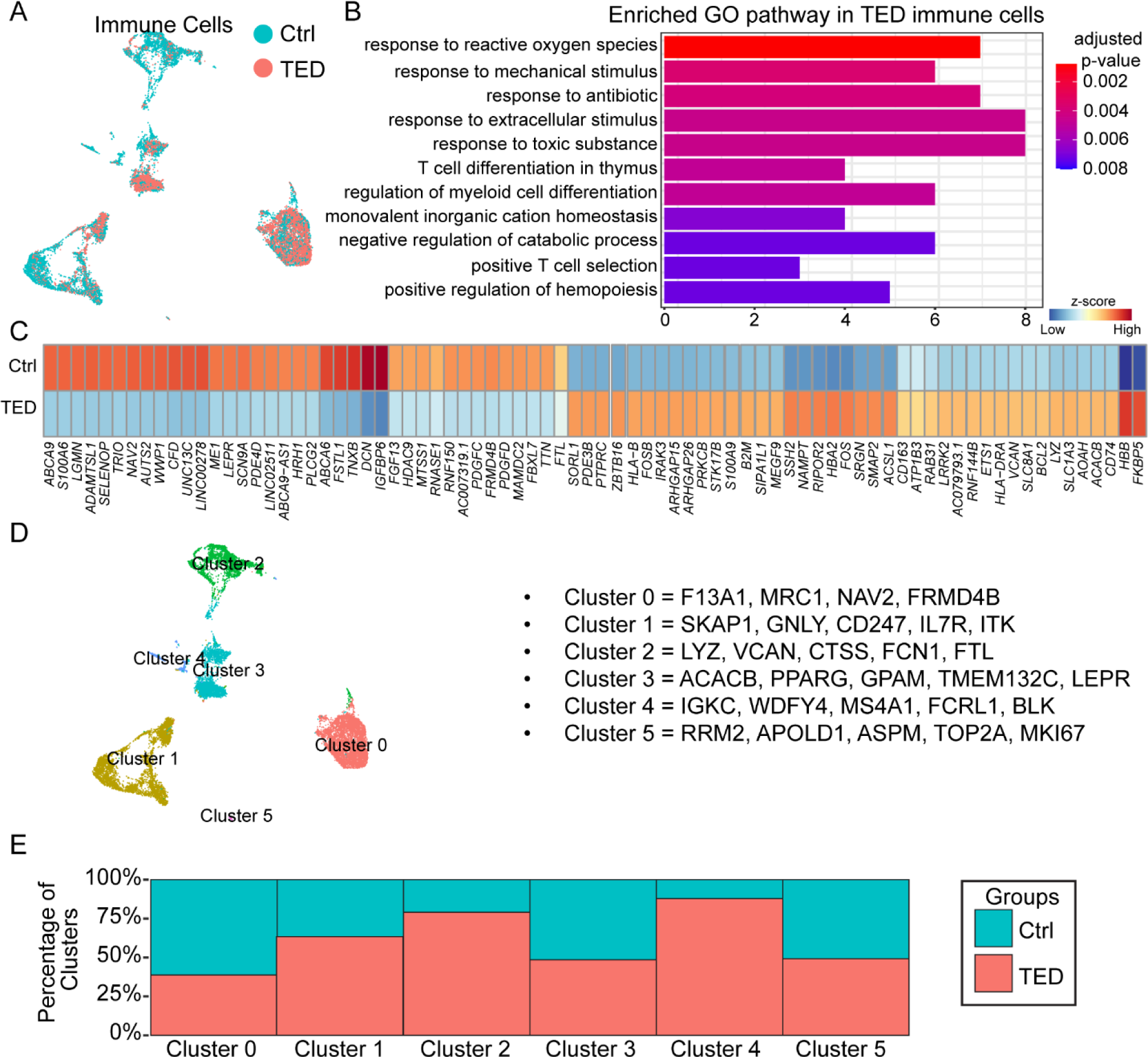
**A.** UMAP plot of immune cells in the control and TED groups. **B.** GO pathway analysis for genes enriched in TED immune cells. **C.** Heatmap displaying the expression of control- or TED-enriched genes in immune cells. **D**. UMAP plot of identified sub-clusters of immune cells in all samples. **E**. Bar graphs comparing the distribution of control and TED groups across the identified clusters in snRNA-Seq.

Further subdivision of immune cells into six distinct clusters (Figure 2D) revealed diverse contributions from each cluster between the groups (Figure 2E). Although the specific immune cell subtypes within each cluster remained unidentified (Table S4), the markers noted suggest a composition enriched in monocytes/macrophages (Clusters 0, 2), NK cells and T cells (Cluster 1), dendritic cells (Clusters 2, 3), and B cells (Cluster 4). Cluster 1 cells from TED patients showed elevated expression of CD4-positive T cell markers, aligning with a previous study that identified CD4 T cells associated with hyperthyroidism and TED (33). These markers include *PRF1*, *GZMA*, *GNLY*, and *GZMH* (Figure S4).

### Differential Expression of the IGF Signaling Pathway Among TED Orbital Fibroblasts

Further investigation of orbital fibroblast populations from TED and control samples identified multiple sub-clusters of orbital fibroblasts, each distinguished by unique molecular markers (Figure 3A, B, Table S5). A GO pathway analysis of genes enriched within these clusters identified varied biological pathways (Figure S5). While some clusters showed no unique pathway alterations, the clusters predominantly found in TED samples exhibited distinctive pathway modifications (Figure S5). Specifically, clusters labeled as ‘NFATC2’, ‘TSHR & PDE10A’, and ‘XACT’ were markedly prevalent in TED samples (Figure 3C), and these clusters displayed changes in pathways related to ‘response to steroid hormone’ and ‘cellular response’. These findings suggest biological shifts associated with TED or impacts of therapeutic interventions like radiotherapy or drug treatments on mitigating TED symptoms (Figure S5).

**Figure 3.**
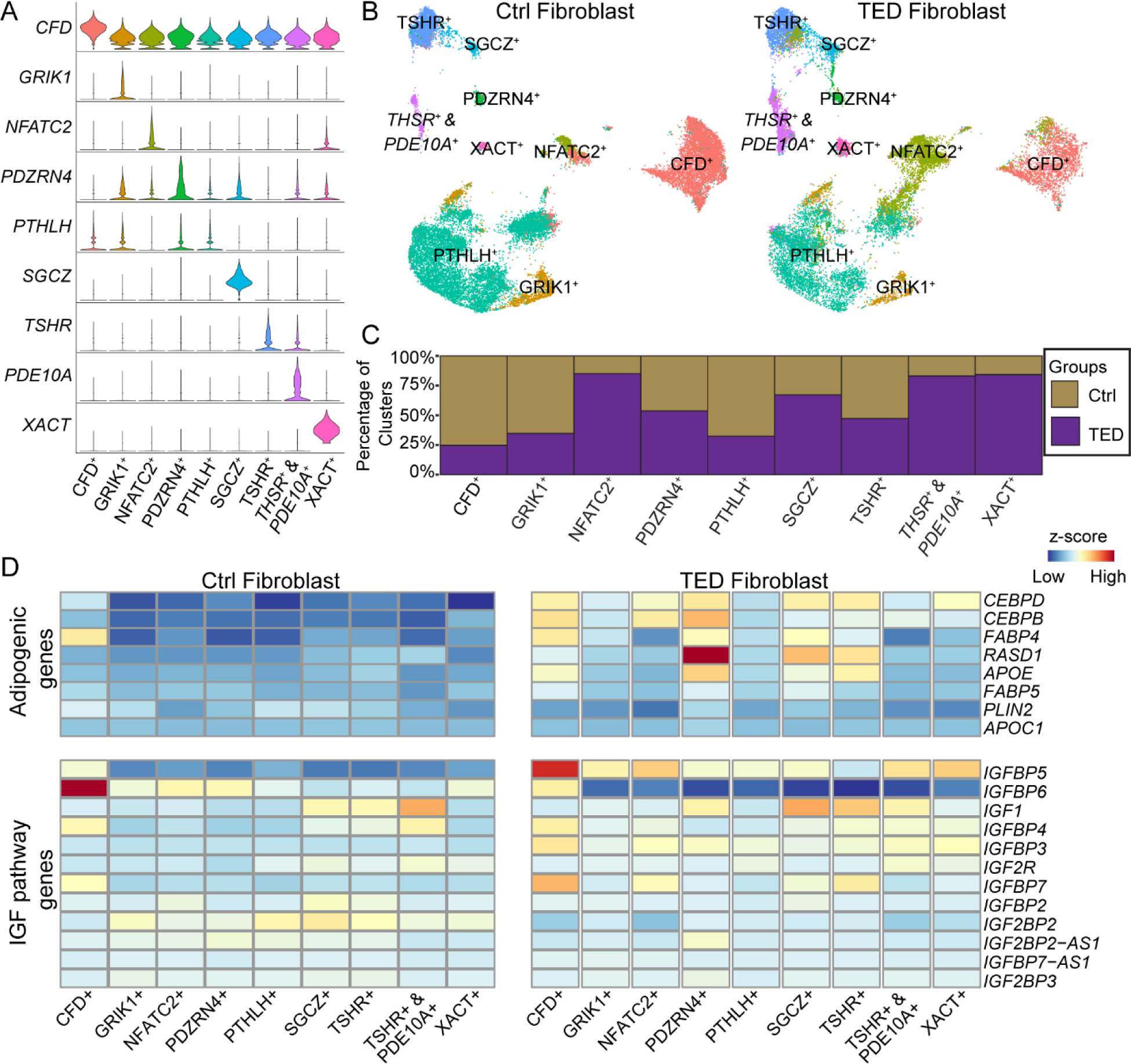
**A.** Violin plots showing genes enriched in specific clusters across orbital fibroblast subtypes in all samples combined. **B.** UMAP plot showing the distribution of orbital fibroblast sub-clusters in control (left) and TED (right). **C.** Bar graphs comparing the distribution of control and TED groups across the orbital fibroblast sub-clusters. **D**. Heatmap displaying the expression of adipogenic and IGF-related genes across groups.

Despite the variability in molecular markers across fibroblast sub-clusters, a commonality in the expression of adipocyte-enriched genes (18) was observed between TED and control groups (Figure S6). TED orbital fibroblasts, irrespective of the cluster, demonstrated higher levels of adipocyte-enriched genes compared to controls (Figure S6). Additionally, an examination of genes known to regulate adipogenesis, such as *CEBPB* and *CEBPD* (34), *FABP4* (*35*), *FABP5* (*36*), *PLIN2* (37), *APOC1* (38), *APOE* (*39*), and *RASD1* (40), revealed a comprehensive upregulation in adipogenic genes, notably *CEBPB* and *CEBPD* in all TED-associated orbital fibroblast sub-clusters (Figure 3D, Table S6). This suggests an inherent adipogenic predisposition within TED fibroblasts. However, other markers like *APOE*, *RASD1*, and *FABP4* were selectively upregulated in specific TED fibroblast sub-clusters (Figure 3D, Table S6). Interestingly, the presence of fibroblast sub-clusters with a higher representation in TED samples did not directly correlate with increased adipogenic gene expression, and no differences were found in the expression of *FABP5, PLIN2,* and *APOC1* between the groups.

Our study also identified disparities in the expression of genes related to the insulin-like growth factor (IGF) signaling pathway across the fibroblast sub-clusters. Notably, TED orbital fibroblasts exhibited a significant reduction in *IGFBP6* expression alongside an increase in *IGFBP5* relative to controls (Figure 3D, Table S6). This differential expression indicates a distinctive modulation of the IGF signaling pathway in TED fibroblasts, suggesting a unique pathophysiological mechanism at play.

### Enhanced Adipogenesis in TED Adipocytes

In our analysis of adipocytes derived from both TED and control groups, we identified multiple sub-clusters within each group, each defined by distinct molecular markers (Figure 4A, Table S7). Notably, the TED group exhibited a significantly increased number of adipocytes compared to the control group (Figure 4B).

**Figure 4.**
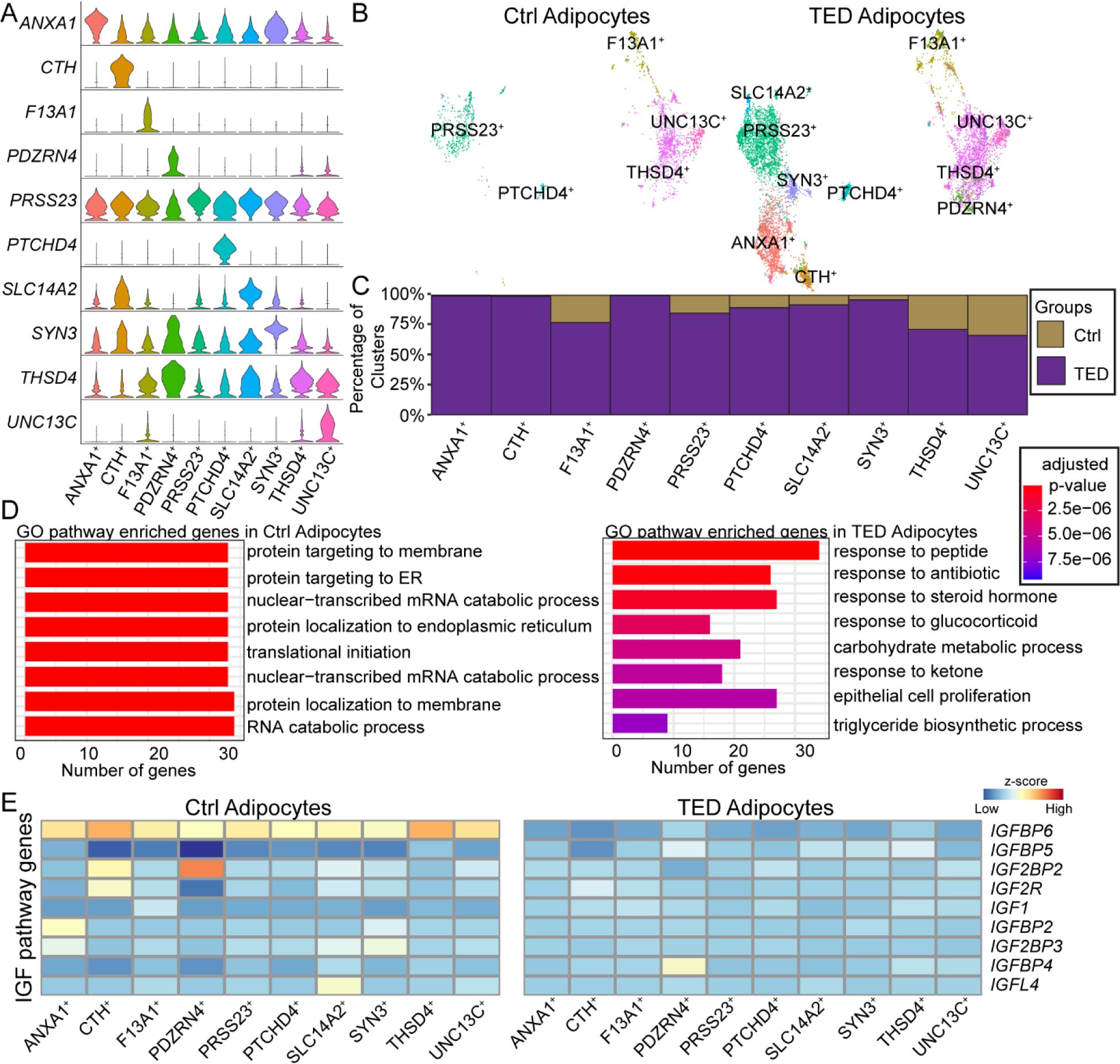
**A.** Violin plots showing genes enriched in specific clusters across orbital adipocyte subtypes in all samples combined. **B.** UMAP plot showing the distribution of orbital adipocyte sub-clusters in control(left) and TED (right). **C.** Bar graphs comparing the distribution of control and TED groups across the orbital adipocyte sub-clusters. **D**. GO pathway analysis of genes enriched in control (left) and TED adipocytes (right). **E**. Heatmap displaying the expression of IGF-related genes across groups.

Additionally, unique adipocyte sub-clusters named ‘ANXA1’, ‘CTH’, and ‘PDZRN4’ were predominantly found in TED samples (Figure 4C). A GO pathway analysis of these sub-clusters, particularly the ‘CTH’ sub-cluster, indicated a distinct shift towards ‘fat cell differentiation’ (Figure S7), suggesting these sub-clusters may represent recently differentiated adipocytes.

Further comparative GO pathway analysis across all sub-clusters of control and TED adipocytes revealed notable differences (Figure 4D, S7). TED adipocytes were enriched in pathways associated with ‘response to peptide’, ‘response to steroid hormone/glucocorticoid’, and ‘proliferation’. These findings suggest a complex interaction of differentiation and proliferation processes contributing to the increased adipocyte population observed in TED (Figure 4D).

A detailed examination of gene expression within the IGF pathway between the two cohorts further uncovered differences. Despite variability within sub-clusters, TED adipocytes consistently displayed a decrease in *IGFBP6* expression across all sub-clusters compared to controls (Figure 4E, Table S8), alongside an increase in *IGFBP5* expression (Table S8). Additionally, reductions in *IGFL4* and *IGFBP2* expression were observed in TED adipocytes, in contrast to the mixed expression patterns of IGF pathway genes seen in orbital fibroblasts. This pattern indicates a more pronounced dampening of IGF signaling in TED adipocytes.

### Adipogenesis Trajectory of TED Orbital Fibroblasts and Adipocytes

To investigate the adipogenic differentiation of orbital fibroblasts in TED, we conducted an integrated analysis of TED orbital fibroblasts with adipocytes. Due to the limited number of adipocytes in control samples, this analysis was specifically focused on identifying changes within the TED cohort.

Our initial objective was to map a direct adipogenic pathway from specific fibroblast sub-clusters to differentiated adipocytes. Although the ‘NFATC2’ fibroblast cluster in TED exhibited a closer association with differentiated adipocytes compared to the ‘CFD’ cluster, a definitive adipogenic lineage from any particular fibroblast sub-cluster to adipocytes could not be conclusively established (Figure 5A).

**Figure 5.**
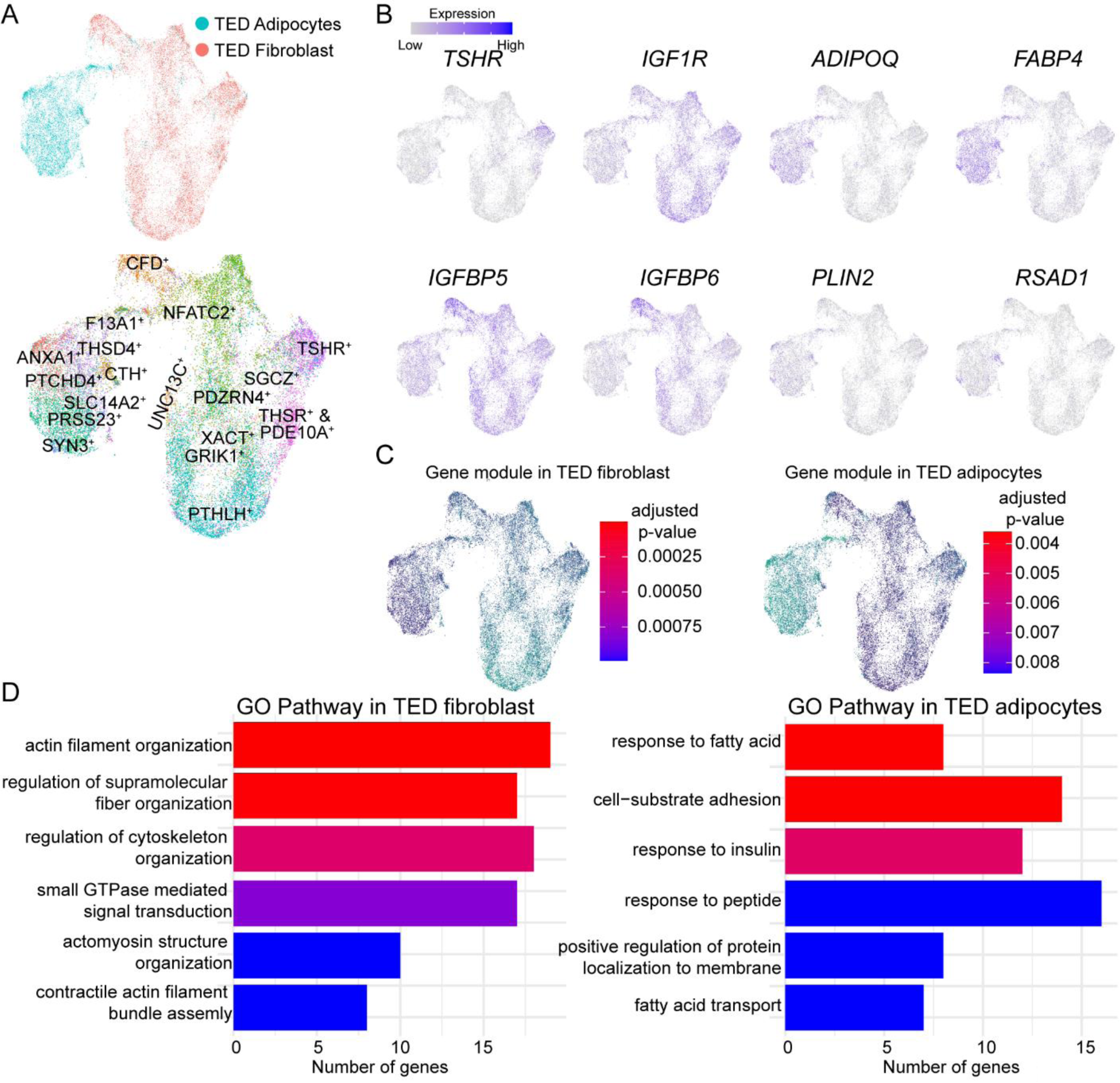
**A.** UMAP plot showing merged orbital fibroblasts and adipocytes in the TED group (top) and previously identified clusters (bottom). **B.** UMAP plots showing the expression of cluster-specific/enriched genes. **C.** Gene modules in TED fibroblast (left) and TED adipocytes (right). **D**. GO pathway analysis of gene modules in TED fibroblasts (left) and TED adipocytes (right).

A significant observation was the downregulation of *TSHR* in adipocytes compared to orbital fibroblasts (Figure 5B), accompanied by a consistent decrease in the expression of *IGF1R, IGFBP5*, and *IGFBP6* (Figure 5B). Conversely, adipocytes demonstrated an upregulation of adipogenic markers such as *ADIPOQ*, *FABP4, PLIN2,* and *RASD1* (Figure 4b), indicating successful adipogenesis.

A GO pathway analysis delineated distinct biological processes between the two cell types (Figure 5C). TED orbital fibroblasts were predominantly involved in pathways associated with ‘cytoskeleton organization/assembly’ (Figure 5D). In contrast, TED adipocytes were linked to ‘response to fatty acid’ and ‘insulin response’ pathways (Figure 5D), underscoring the significant role of insulin in adipogenic differentiation from fibroblast to adipocytes (18). These observations further reinforce the importance of the IGF pathway in the adipogenic process in TED, aligning with findings from earlier *in vitro* studies on patient-derived human orbital fibroblast cell lines (18).

### Linsitinib Treatment Reduces Adipogenesis in TED Fibroblasts

The observed decrease in IGF-1 pathway gene expression in differentiated adipocytes relative to orbital fibroblasts aligns with previous research that IGF-1 signaling initially surges during the early stage of adipogenesis *in vitro*, then declines as adipocytes mature. This pattern prompted us to investigate the effects of manipulating the IGF-1 pathway on adipogenesis in orbital fibroblasts. We utilized linsitinib, a small molecule IGF-1R antagonist, which diminishes immune infiltration and fibrosis in the orbit in TED animal models (41). Currently, linsitinib is undergoing Phase 2 clinical trials as a potential oral therapy for TED.

In our established *in vitro* model using patient-derived human orbital fibroblast cell lines (18), we administered linsitinib in a dose-dependent manner (Figure 6A-L). Consistent with prior observations, adipocyte differentiation was evident by Day 5 *in vitro*, as indicated by Oil Red O-positive adipocytes (Figure 6E). A notable dose-dependent reduction in the proportion of adipocytes was documented following linsitinib treatment on both Days 5 and 9 (Figure 6M). Considering linsitinib’s known affinity for the insulin receptor (42), which also plays a role in adipogenesis, we conducted additional experiments in insulin-free media to show the insulin-independent effects of IGF-1R inhibition on adipogenesis. Despite the overall reduction in adipogenesis due to the absence of insulin, a critical factor in adipocyte differentiation, linsitinib treatment still resulted in a dose-dependent inhibition of adipogenesis (Figure 6N). To confirm that the decrease in adipogenesis associated with linsitinib treatment is due to inhibition of IGF-1R signaling, we performed Western blots to demonstrate that treatment with linsitinib is associated with a dose-dependent decrease in IGF-1R phosphorylation on days 5 and 9 (Figure 6O). These findings emphasize the critical role of IGF-1R signaling in the adipogenic process of TED orbital fibroblasts, highlighting how linsitinib’s influence on this pathway significantly impacts adipogenesis, independent of insulin signaling.

**Figure 6.**
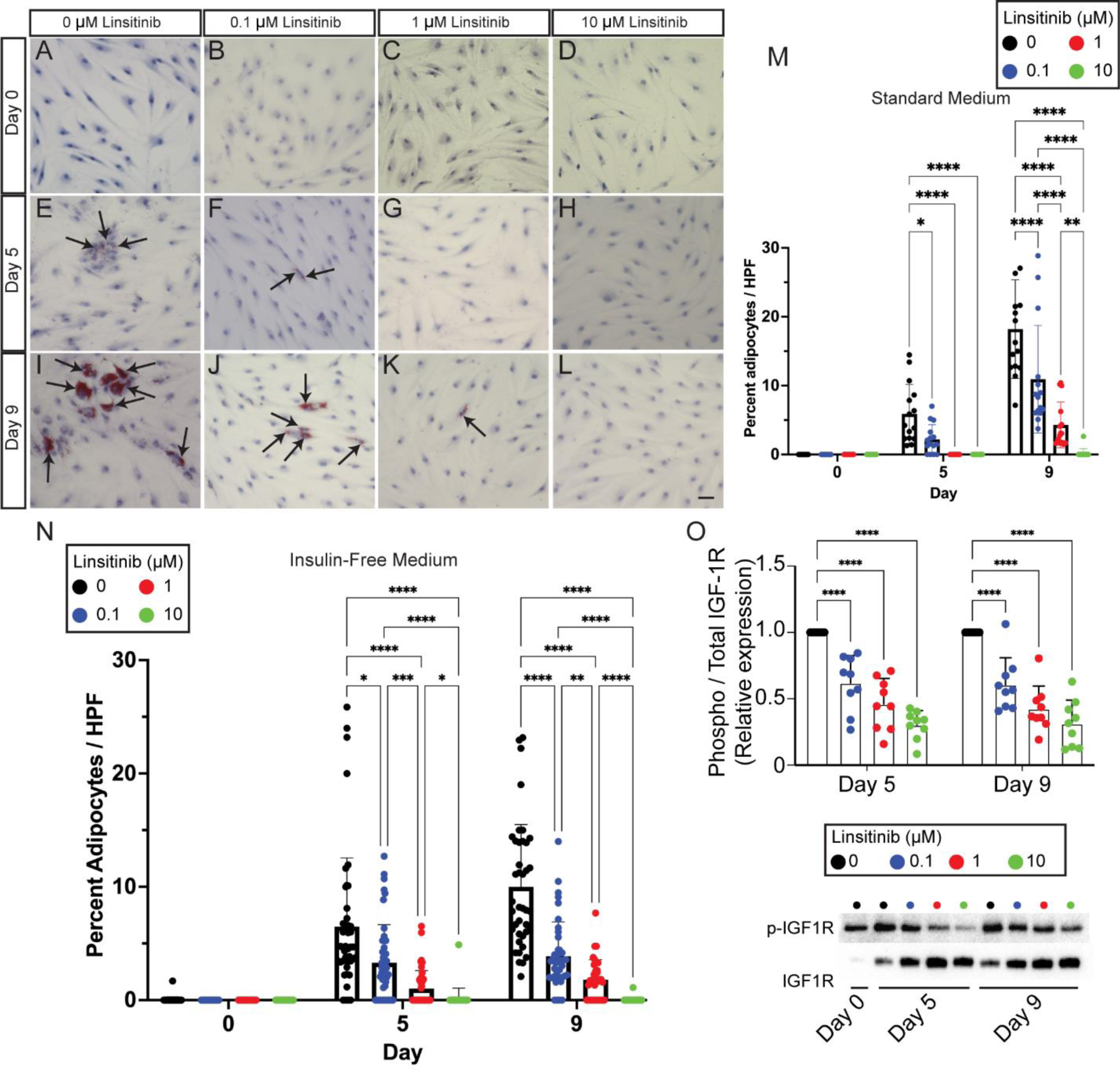
**A-L.** Oil Red O staining of orbital fibroblasts treated with adipogenic media (A, E, I) and 0.1 μM (B, F, J), 1 μM (C, G, K), or 10 μM (D, H, L) linsitinib days 0 (A-D), 5 (E-H) and 9 (I-L) after treatment. Arrows indicate Oil Red O positive adipocytes. Scale bar = 50 μm. **M.** Bar plot showing the percentage of adipocytes seen after treatment with standard adipogenic medium and 0, 0.1, 1, or 10 uM linsitinib on days 0, 5, and 9. **N.** Bar plot showing the percentage of adipocytes seen after treatment with insulin-free adipogenic medium and 0, 0.1, 1, or 10 uM linsitinib on days 0, 5, and 9. **O.** Bar plot (top) and blots (bottom) showing the relative expression between phospho-IGF1R and total IGF1R after treatment with insulin-free adipogenic medium and 0, 0.1, 1, or 10 uM linsitinib on days 5 and 9. * indicates *p < 0.05*; ** indicates *p < 0.01*; *** indicates *p < 0.001*; **** indicates *p < 0.0001*.

## Discussion

In this study, we utilized snRNA-Seq to explore the cellular landscape of orbital fat in TED, providing the first *in vivo* evidence of increased adipogenesis linked to this condition. We observed a significant rise in the number of adipocytes within the orbital fat of TED patients, alongside differential expression of components in the IGF-1 signaling pathway. Specifically, orbital fibroblasts in TED patients were enriched with IGF-1 pathway components, contrasting with the decreased pathway activity observed in TED adipocytes. These observations extend our previous *in vitro* work (18), reinforcing the hypothesis that disrupted IGF-1 signaling is crucial in TED pathogenesis.

Highlighting the complexity of IGF signaling dysfunction in TED, we found an upregulation of *IGFBP5* and downregulation of *IGFBP6* in TED orbital fibroblasts, which diminished as the fibroblasts differentiated into adipocytes (Figure 5B). This is consistent with previous work indicating that IGFBP5 is upregulated alongside IGF signaling in vascular smooth muscle cells (43).

Additionally, we found that TED immune cells exhibit higher levels of *IGFBP6* expression compared to those from controls, indicating that IGF1 pathway dysfunction in the orbit is not limited to fibroblasts and adipocytes (Table S3). The role of IGFBP5 in modulating IGF signaling is complex, potentially inhibiting (44, 45) or enhancing (46, 47) IGF signaling, depending on calcium concentrations in the surrounding environment (48). Both IGFBP5 and IGFBP6 bind IGF2 more strongly than IGF1 and possess IGF-independent functions, including effects on apoptosis, cell adhesion, and angiogenesis (49–51). Further studies are needed to better describe the levels of IGF1 and IGFBPs in orbital fat to delineate how IGFBP expression modulates IGF1 signaling in TED. Moreover, investigating the potential IGF-independent roles of IGFBPs in orbital fibroblasts will be crucial for a more comprehensive understanding of their impact on the disease’s pathophysiology.

Our analyses also highlight two potential mechanisms driving adipocyte accumulation in TED: enhanced differentiation of orbital fibroblasts into adipocytes and increased proliferation of pre-existing adipocytes. Notably, TED adipocytes exhibited a downregulation in the expression of several IGF pathway genes such as *IGFBP5, IGFBP6, IGFL4*, *IGFBP2,* and *IGF1R* relative to fibroblasts. This reduced IGF signaling is likely a key factor contributing to TED progression and indicates a shift in cellular signaling pathways that warrants further exploration.

In TED orbital fibroblasts, we observed not only the upregulation of adipocytes-enriched genes but also higher expression levels of genes that regulate adipogenesis, including *CEBPB* and *CEBPD* (34), *FABP4* (35), *RASD1* (40), and *APOE* (39). However, we detected no changes in other adipogenic genes such as *FABP5* (36), *PLIN2* (37), and *APOC1* (38). The specific roles of these genes in adipogenesis require further investigation. Additionally, we identified other genes in TED fibroblasts that may facilitate adipogenesis (Table S8), such as *NFATC2,* which can regulate phospholipase A2 (52), a potential adipogenesis target (53), and induces adipocytes in 3T3 cells (54, 55). ZBTB16 also induces adipogenesis *in vitro* (56) and is recognized as a potential adipogenic factor (57). Furthermore, silencing *LRP1B* in rodents has been demonstrated to inhibit adipogenesis (58).

The use of snRNA-Seq in our study offers a refined approach for profiling the cellular composition of orbital fat in TED. This choice is particularly advantageous over single-cell RNA-Seq (scRNA-Seq) due to its enhanced capacity for capturing adipocyte populations, which are typically challenging due to their physical properties (59–61). Interestingly, our results do not fully align with scRNA-Seq data previously published by Li and colleagues (62), where adipocytes were not identified, a discrepancy likely due to selective loss of these cells. Furthermore, the samples profiled by Li and colleagues were isolated from patients with generally higher CASs and greater severity requiring decompression for compressive optic neuropathy, which may account for the differences in the distribution of immune cells described (62).

In this snRNA-Seq analysis, we did not detect SMA-positive myofibroblasts, which are considered critical for fibrosis in TED (63–67). These cells have been described *in vitro* following the treatment of orbital fibroblasts with TGF-beta (67, 68). The absence in our samples could be attributed to the relatively low CASs of the sampled patients, suggesting a correlation between lower CASs and less active fibrosis due to the absence of this cell population. Alternatively, the lack of SMA+ myofibroblasts might be a sampling artifact, as these cells may not reside in the orbital fat but rather in other orbital tissues like extraocular muscles, Tenon’s fascia, or muscle sheaths, which are not routinely biopsied in TED. Profiling additional orbital fat samples from patients at various disease stages and activity levels may provide further insights into the fibroblast populations mediating fibrosis in TED. This absence might also indicate that *in vivo* conversion of preadipocyte fibroblasts into SMA+ myofibroblasts does not occur in patients with a CAS less than or equal to 3.

While tissue culture models have significantly advanced our understanding of TED by allowing for manipulations of cell signaling pathways to uncover disease mechanisms, the use of serially passaged cells has inherent limitations. Our study’s snRNA-Seq data provide additional validation for the role of adipogenesis in TED and suggest the therapeutic potential of IGF-1R inhibition. However, the cross-sectional design of our study limits our ability to collect longitudinal data, which could offer deeper insights into the disease’s progression. Additionally, our dataset involves a limited number of patients with relatively low CASs, all presenting for surgery during the study period. Future studies should aim to include a broader patient cohort, encompassing various stages of TED, to more comprehensively elucidate the progression and cellular dynamics within the orbit.

In particular, further studies of the role of immune cells in the progression of TED, the potential interplay between inflammation, adipogenesis, and adipocyte proliferation on TED pathology, and the impact of thyroid hormone regulation of IGF-1 signaling versus treatment effects should be undertaken.

## Acknowledgments

This work was supported by a Research to Prevent Blindness Physician Scientist Award to FR, an unrestricted departmental grant to the Wilmer Eye Institute from Research to Prevent Blindness, the Johns Hopkins University School of Medicine Core Coins Program, and Lundbeckfonden grant (R361-2020-2654) to DWK.

## Author contributions

Designing research studies: FR, Conducting experiments: DWK, SK, JH, KB; Analyzing data: DWK, SK, FR; Providing reagents: EL, NM, SB, FR; Funding: DWK, SB, FR; Writing the manuscript: DWK, FR; Critical review of the manuscript: DWK, SB, FR.

---

**Figure S1.**
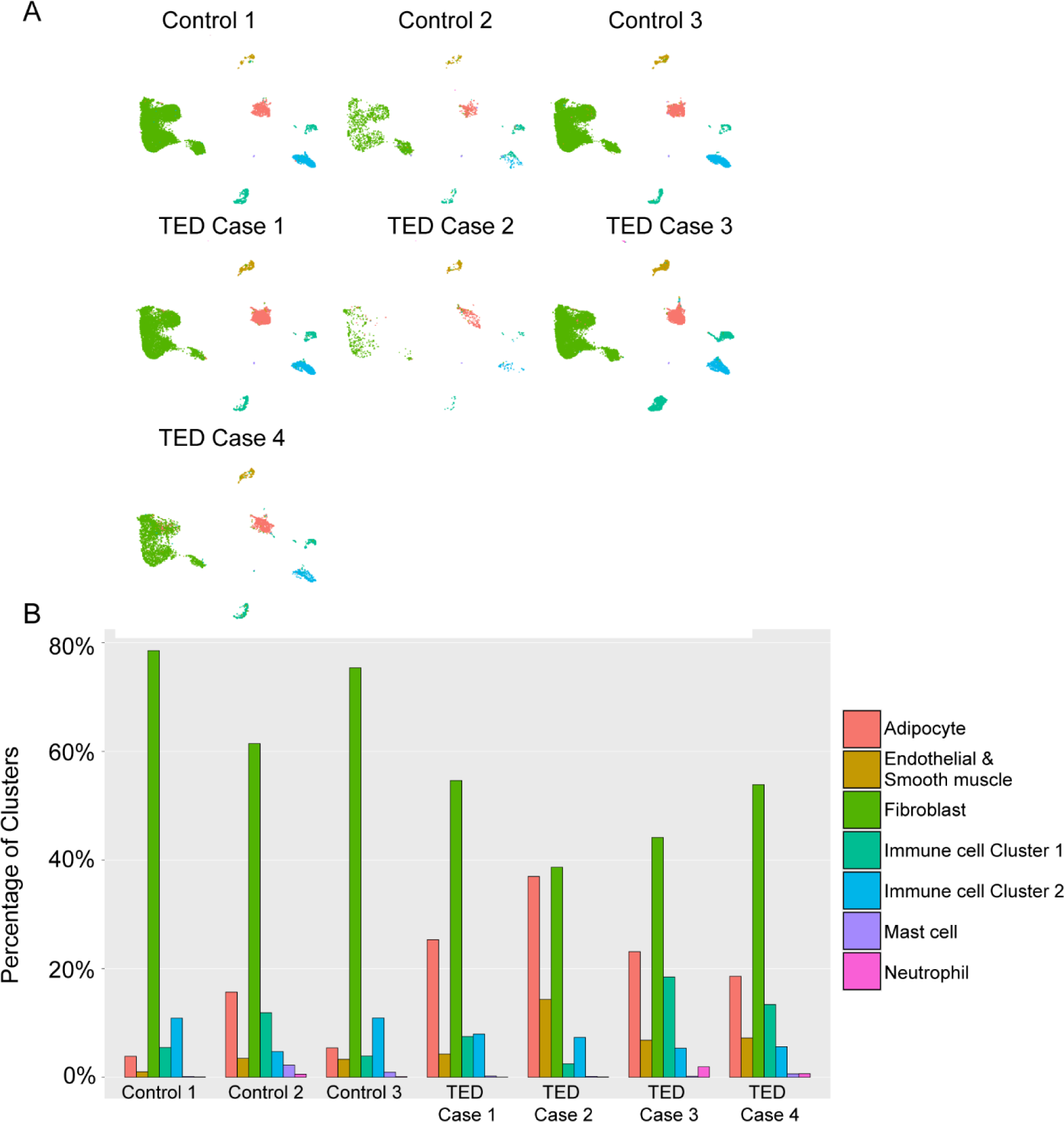
**A.** UMAP plot illustrating the distribution of cell types for each sample. **B.** Bar graphs showing the distribution of each cell type in each sample.

**Figure S2.**
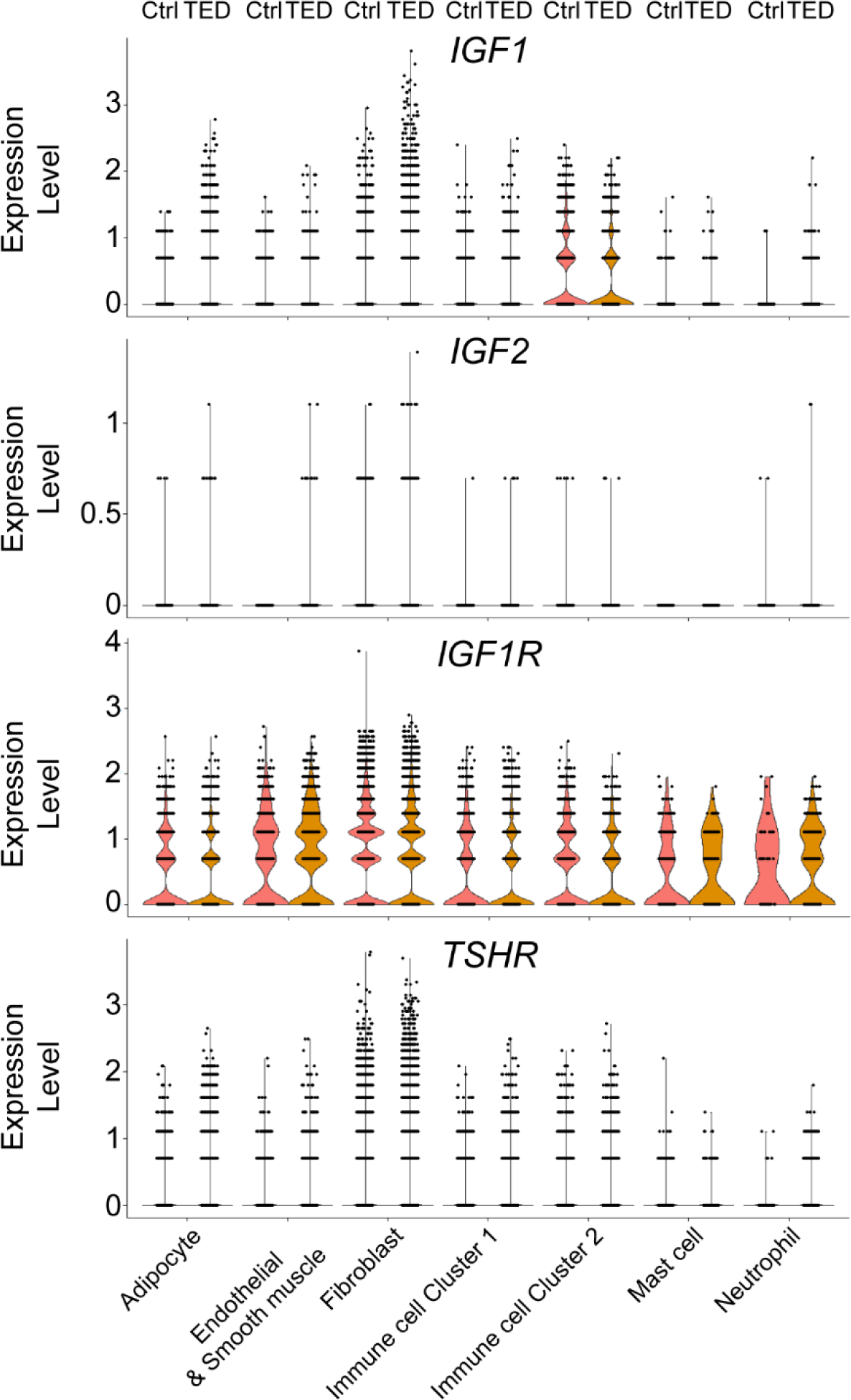
Violin plots showing the expression levels of *IGF1*, *IGF2*, *IGF1R*, and *TSHR* across various cell types in both control and TED groups.

**Figure S3.**
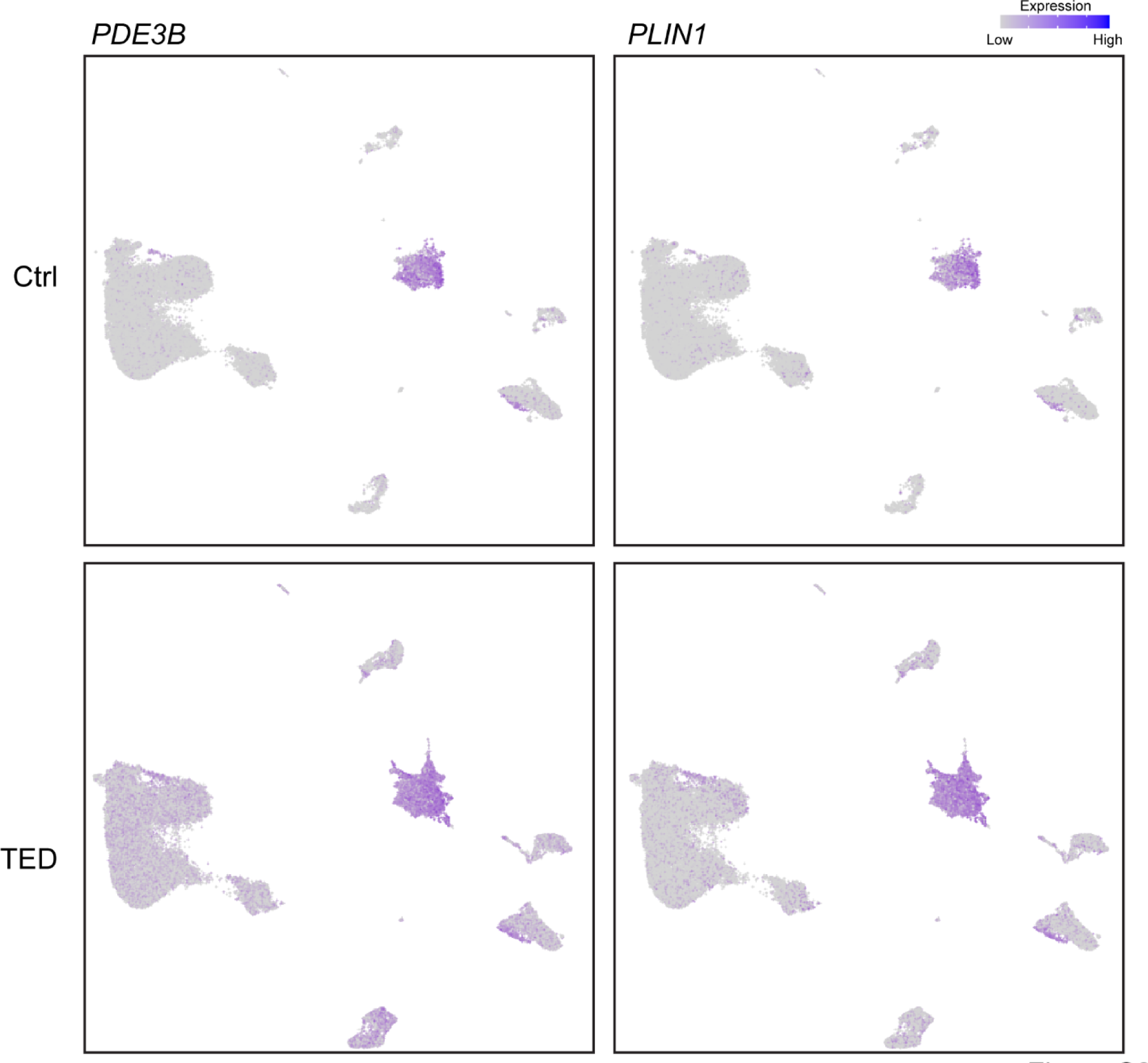
UMAP plot illustrating the expression of *PDE3B* and *PLIN1* in each cell type across control and TED groups.

**Figure S4.**
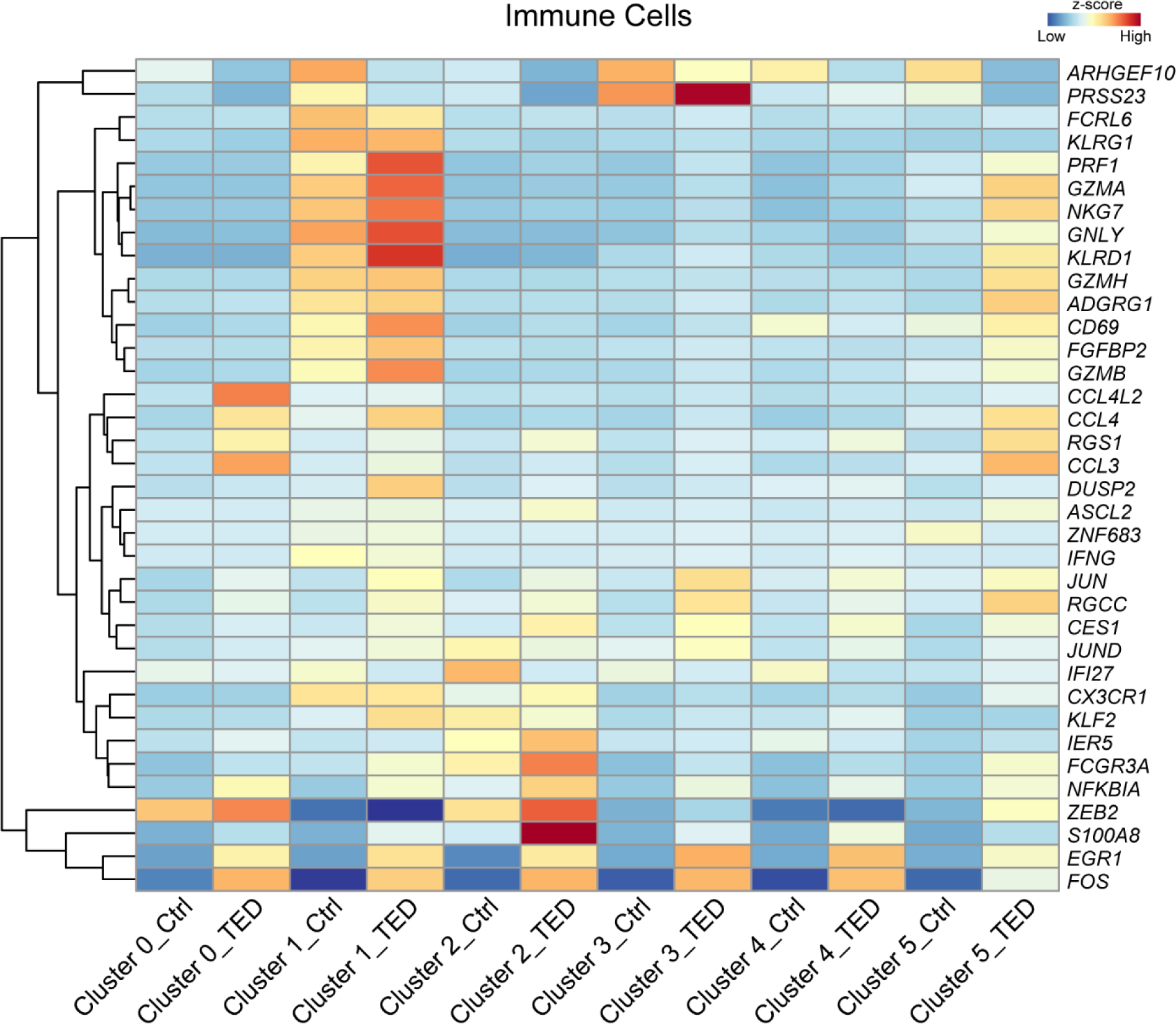
Heatmap showing the expression patterns of TED-associated CD4 T cell markers (33).

**Figure S5.**
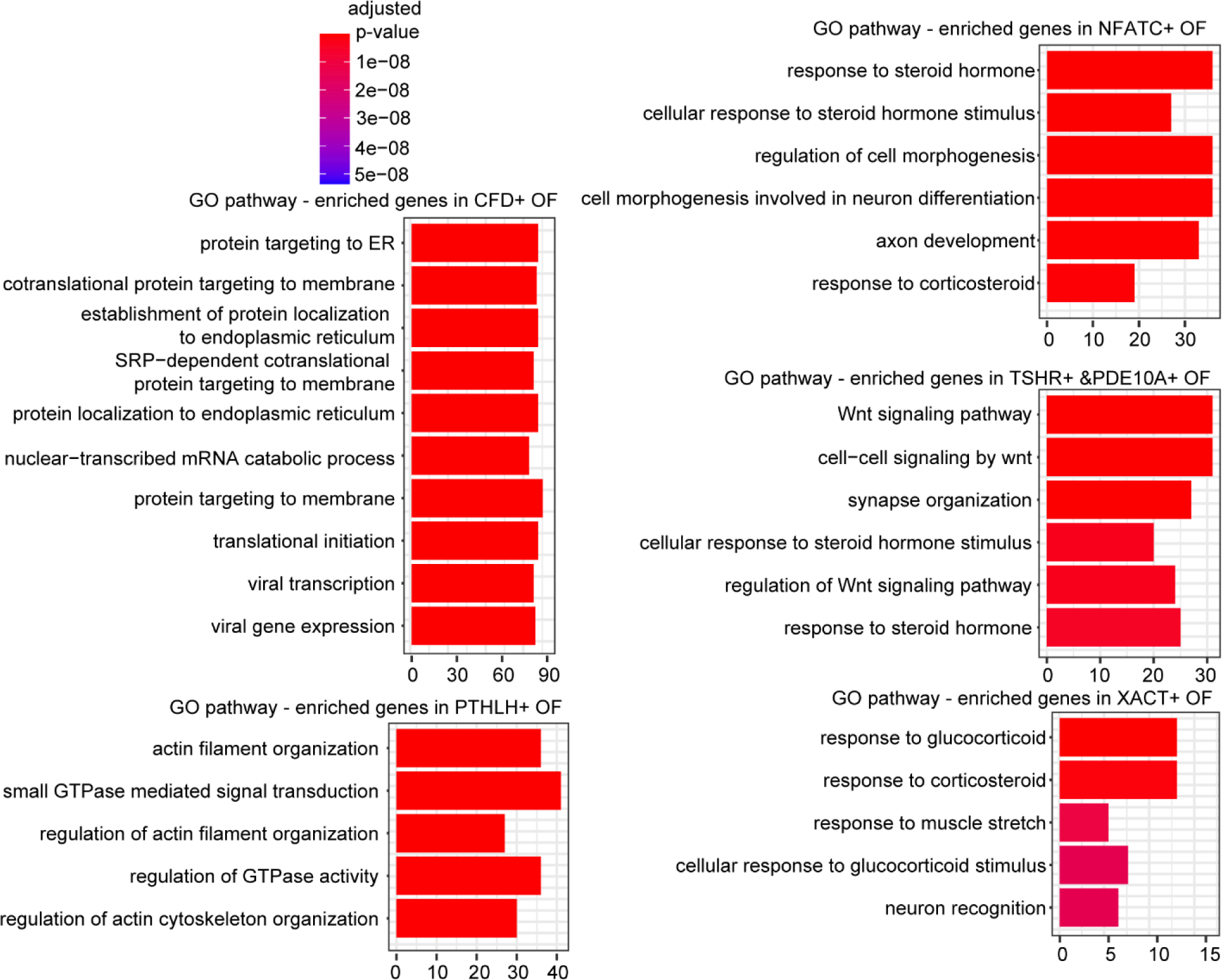
GO pathway analysis for specific pathways displayed across different subtypes of orbital fibroblasts.

**Figure S6.**
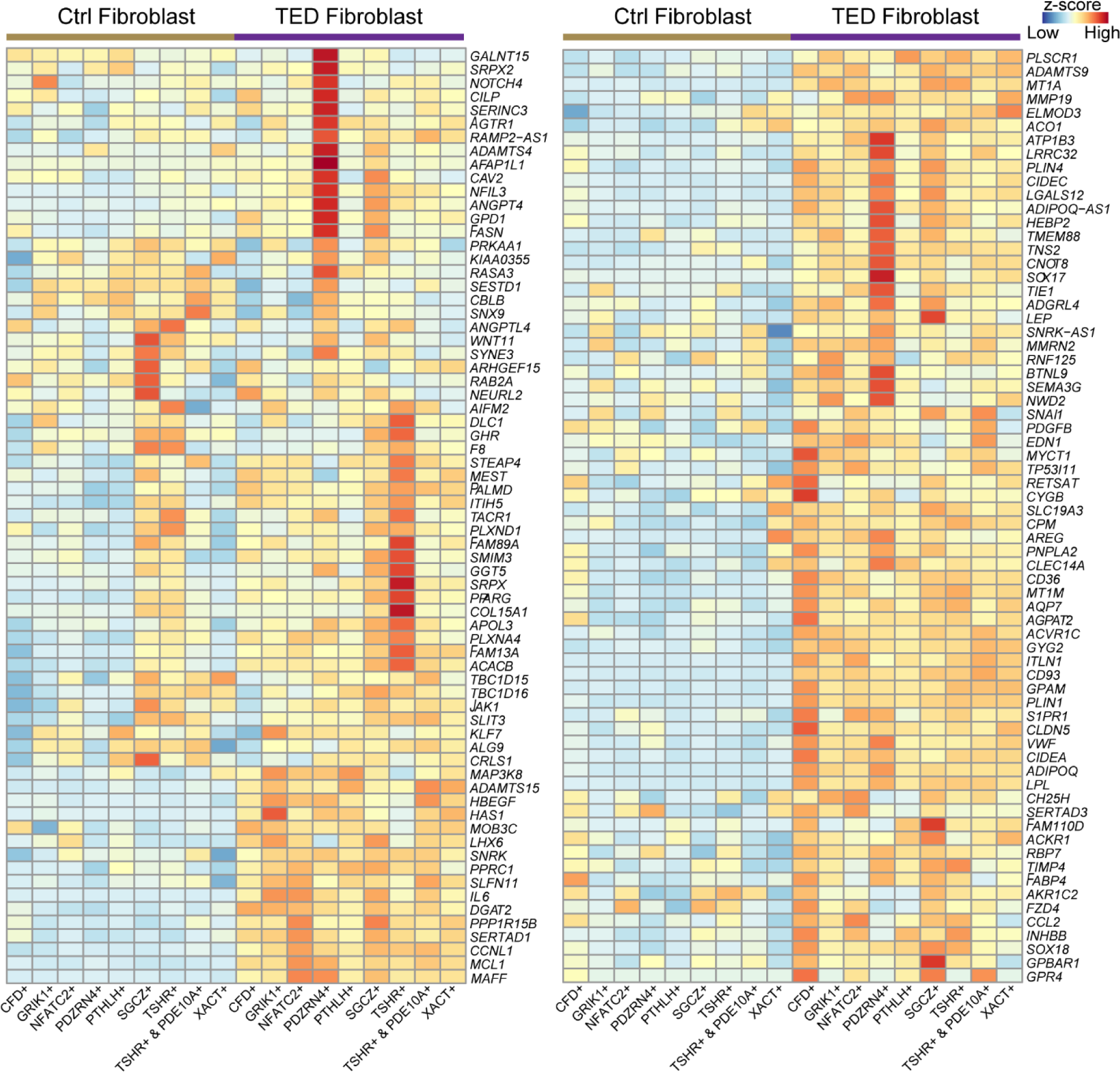
Heatmap illustrating the distribution of adipocyte-enriched genes across control and TED orbital fibroblast subtypes.

**Figure S7.**
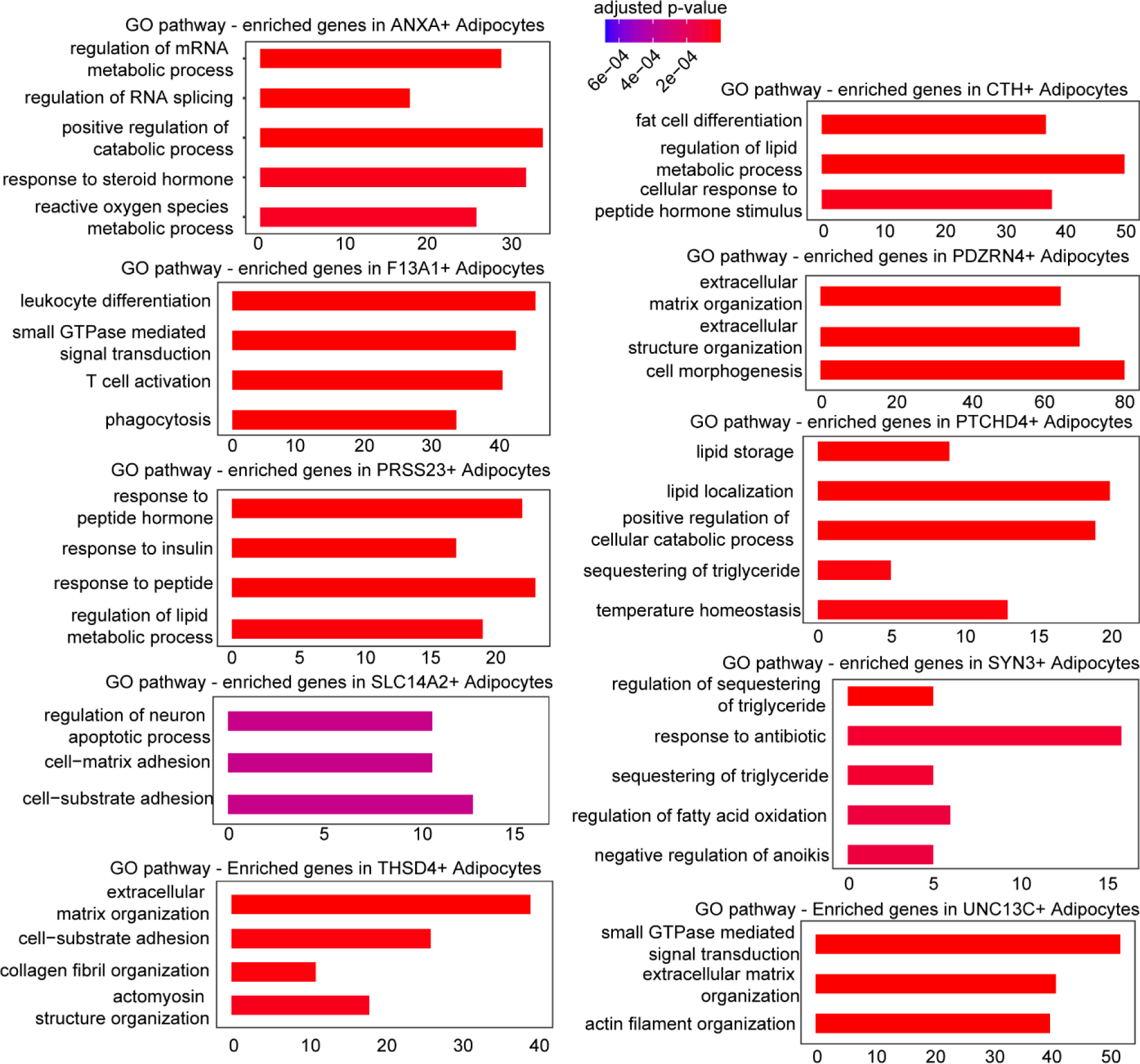
GO pathway analysis for specific pathways displayed across different subtypes of orbital adipocytes.

## Table Legends

Table S1. Summary of snRNA-Seq samples utilized in this study.

Table S2. Analysis of differential gene expression across identified clusters in snRNA-Seq data.

Table S3. Comparative analysis of differential gene expression in immune cells between control and TED groups.

Table S4. Differential gene expression analysis within immune cell sub-clusters.

Table S5. Differential gene expression analysis within orbital fibroblast sub-clusters.

Table S6. Comparison of differential gene expression in orbital fibroblasts between control and TED groups.

Table S7. Differential gene expression analysis within orbital adipocyte sub-clusters.

Table S8. Comparison of differential gene expression in orbital adipocytes between control and TED groups.

